# Differentially expressed gene profile and relevant pathways of the inhibition of LAMP5 on MLL-rearranged leukemias

**DOI:** 10.1101/2022.08.17.504277

**Authors:** Zhiwen Qian, Wei Zhang

**Affiliations:** Shanghai University of Medicine and Health Sciences

## Abstract

LAMP5 is a lysosome-related protein, which showed increased expression levels in MLL-r leukemia. Recent findings showed that LAMP5 is involved in MLL-fusion protein but the specific mechanism is still unclear. In our study, we aim to identify the transcriptional landscapes of the LAMP5-shRNA-treated MLL-r leukemia cells. We used the gene enrichment analyses such as KEGG and GO to further analyze the potential signaling pathways. Moreover, we constructed the PPI network and Reactome map to further identify the biological processes. We found that Lysosome and Cell Cycle are the major signaling pathways involved in the LAMP5-shRNA treated MLL-r leukemia cells. We identified the top ten interactive genes including CDK1, TOP2A, PLK1, CCNA2, ASPM, BUB1, BUB1B, CCNB1, AURKA, and KIF11. Our study may provide novel mechanisms for the treatment of MLL-r leukemia by inhibiting LAMP5.

## Introduction

The abnormal activity of the MLL gene is the cause of aggressive leukemias and shows in about 10% of human acute leukemias^1^. MLL-leukemia is a kind of acute myeloid leukemia (AML) and acute lymphoblastic leukemia (ALL) that are identified in both children and adults^2^. MLL rearrangement is a poor prognosis maker that is identified in infants with ALL^3^. There is an urgent need for more effective, less toxic therapies. MLL rearrangements create a chimeric gene that encodes a protein that can cause cancers^4^. MLL fusion proteins bind to DNA/chromatin and lead to leukemic transformation in hematopoietic stem and progenitor cells through transcriptional regulation^5^. The best-studied target genes contain HOXA genes, which have been shown to be critical for leukemia development and progression^6^. Other important members of the MLL fusion-driven gene expression program contain CDK6 and MEF2C. Other work has focused on the mechanisms that would lead to new therapeutic approaches^7^.

LAMP-5 belongs to the lysosome-associated membrane protein (LAMP) family, which shows widespread expression, and the Lamp5 expression in mice is confined to several regions of the postnatal brain^8^. LAMP5 is typically expressed in plasmacytoid dendritic cells (pDC) in human brains^9^. LAMP-5 helps the transport of TLR9 from early endosomal to lysosomal signaling vesicles, thereby mediating type 1 interferon and pro-inflammatory signaling respectively^10^. Moreover, LAMP-5 acts as a cell surface protein and plays an essential role in MLLr leukemias through the regulation of innate immune signaling^4^.

In this study, our aim is to identify the role of LAMP-5 in MLL-rearranged acute leukemia by analyzing the RNA-seq data. We further analyzed the KEGG, GO, Reactome map, and PPI for exploring the mechanisms of the knockdown of LAMP-5 in acute leukemia cells. Therefore, our study may provide directions for identifying the drug targets of acute leukemia.

## Methods

### Acquisition of microarray data

GEO database (http://www.ncbi.nlm.nih.gov/geo/) was used for downloading the following gene expression profile dataset: GSE166523. The data was produced by the Illumina HiSeq 2500 (Homo sapiens) (Cincinnati Children’s Hospital, Biomedical Informatics, Nathan Salomonis, 3333 Burnet Avenue, Cincinnati, OH 45229, USA). The dataset was chosen based on the criteria as follows: Studies comparing control shRNAs treated MLL leukemia cells with LAMP-5 shRNAs treated MLL leukemia cells; Studies included gene expression profiles; Datasets were excluded if the study did not contain control groups; Datasets from other organisms were excluded.

### Identification of differentially expressed genes (DEGs)

The gene expression matrix data in the GSE166523 dataset was normalized and processed through the R program as described. Then, the empirical Bayesian algorithm in the “limma” package was applied and compared between patients with diseases and the healthy control population as controls with the detection thresholds of adjusted P-values <0.05 and |log2FC| >1.

### Gene intersection between DEGs

We intersected the DEGs screened by the “limma” package with the modular genes. The intersecting genes were analyzed for Kyoto Encyclopedia of Genes and Genomes (KEGG) and Gene Ontology (GO) enrichment using the “clusterprofiler” package in the R program. The results were then visualized using the “ggplot2” package.

### Protein-protein interaction (PPI) networks

The interacting genes were constructed using Protein-Protein Interaction Networks Functional Enrichment Analysis (STRING; http://string.db.org). The Molecular Complex Detection (MCODE) was used to analyze the PPI networks. This method predicts the PPI network and provides an insight into the biological mechanisms. The biological processes analyses were further performed by using Reactome (https://reactome.org/), and P<0.05 was considered significant.

## Results

### DEGs in MLL-rearranged acute leukemia treated with LAMP-5 shRNAs

To determine the effects of LAMP-5 on MLL-rearranged acute leukemia, we used the RNA-seq data from the GEO database (GSE166523). We identified the 1130 significantly changed genes [DEGs, differentially expressed genes (DEGs), P < 0.01]. The top increased and decreased genes were also indicated by the heatmap (Figure 1). We further presented the top ten significant genes in Table 1.

**Table 1.**

| **Table 1** |  |  |  |
| --- | --- | --- | --- |
| **Entrez gene** | **Gene symbol** | **Fold-change** | **Regulation** |
| **Top 10 down-regulated DEGs** |  |  |  |
| 109286553 | ARHGAP27P1-BPTFP1-KPNA2P3 | -7.946075635 | Down |
| 2550 | GABBR1 | -3.684443557 | Down |
| 721 | C4B | -3.638234913 | Down |
| 166929 | SGMS2 | -3.552841349 | Down |
| 3680 | ITGA9 | -3.502870969 | Down |
| 389136 | VGLL3 | -3.310252783 | Down |
| 729156 | GTF2IRD1P1 | -3.285008187 | Down |
| 110384692 | LOC110384692 | -3.259894772 | Down |
| 3304 | HSPA1B | -3.248198068 | Down |
| 2348 | FOLR1 | -3.176667051 | Down |
| **Top 10 up-regulated DEGs** |  |  |  |
| 2201 | FBN2 | 3.315475628 | up |
| 1756 | DMD | 3.053808223 | up |
| 55638 | SYBU | 3.034450873 | up |
| 4969 | OGN | 2.869198695 | up |
| 150297 | C22orf42 | 2.600728621 | up |
| 222389 | BEND7 | 2.588555017 | up |
| 3291 | HSD11B2 | 2.540267099 | up |
| 126326 | GIPC3 | 2.511293603 | up |
| 439936 | LINC02899 | 2.501121439 | up |
| 101929777 | LOC101929777 | 2.369186806 | up |

**Figure 1.**
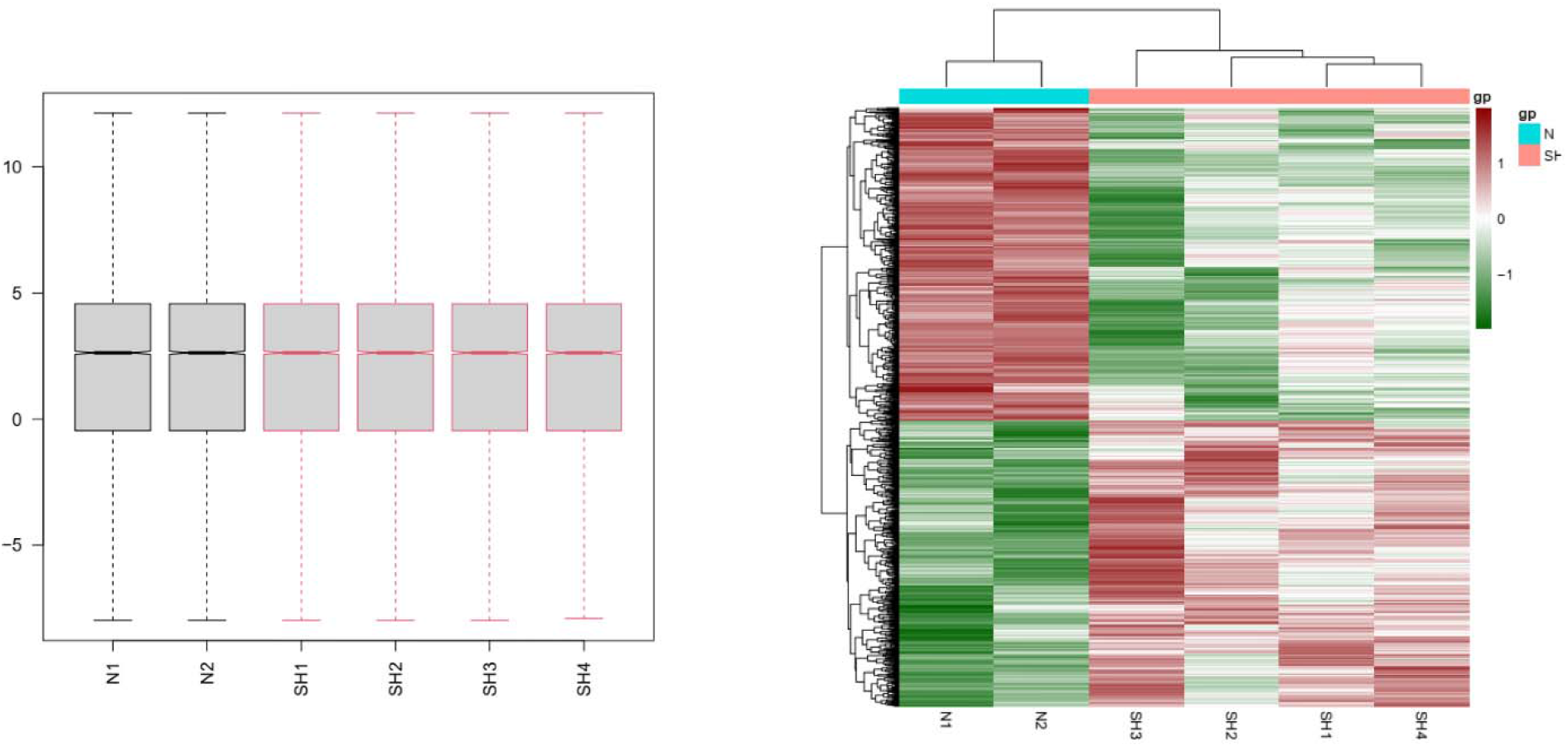
Heatmap in MLL-rearranged acute leukemia treated with LAMP-5 shRNAs. (A) DEGs (P < 0.05) were calculated and used to construct the heatmap. N, control shRNAs; SH, LAMP-5 shRNAs.

### Gene enrichment and function analyses in MLL-rearranged acute leukemia treated with LAMP-5 shRNAs

To determine the features of significant genes, we performed the gene enrichment analysis (Figure 2). We found the top 10 changed KEGG items (P < 0.05), including “Lysosome”, “Cell cycle”, “Nucleocytoplasmic transport”, “Progesterone− mediated oocyte maturation”, “DNA replication”, “Cholesterol metabolism”, “Mismatch repair”, “Base excision repair”, “Homologous recombination”, and “Galactose metabolism”. We found the top 10 changed Biological Processes (BP, P < 0.05), including “organelle fission”, “nuclear division”, “chromosome segregation”, “mitotic nuclear division”, “nuclear chromosome segregation”, “sister chromatid segregation”, “DNA replication”, “mitotic sister chromatid segregation”, “spindle organization”, “mitotic spindle organization”. We identified the top 10 changed Cellular Components (CC, P < 0.05), including “chromosomal region”, “spindle”, “chromosome, centromeric region”, “condensed chromosome”, “condensed chromosome, centromeric region”, “kinetochore”, “mitotic spindle”, “spindle pole”, “CMG complex”, “DNA replication preinitiation complex”. We figured out the top 10 changed Molecular Functions (MF, P < 0.05), including “tubulin binding”, “microtubule binding”, “catalytic activity, acting on DNA”, “microtubule motor activity”, “DNA helicase activity”, “single− stranded DNA helicase activity”, “nuclear localization sequence binding”, “DNA replication origin binding”, “lamin binding”, “histone kinase activity”.

**Figure 2.**
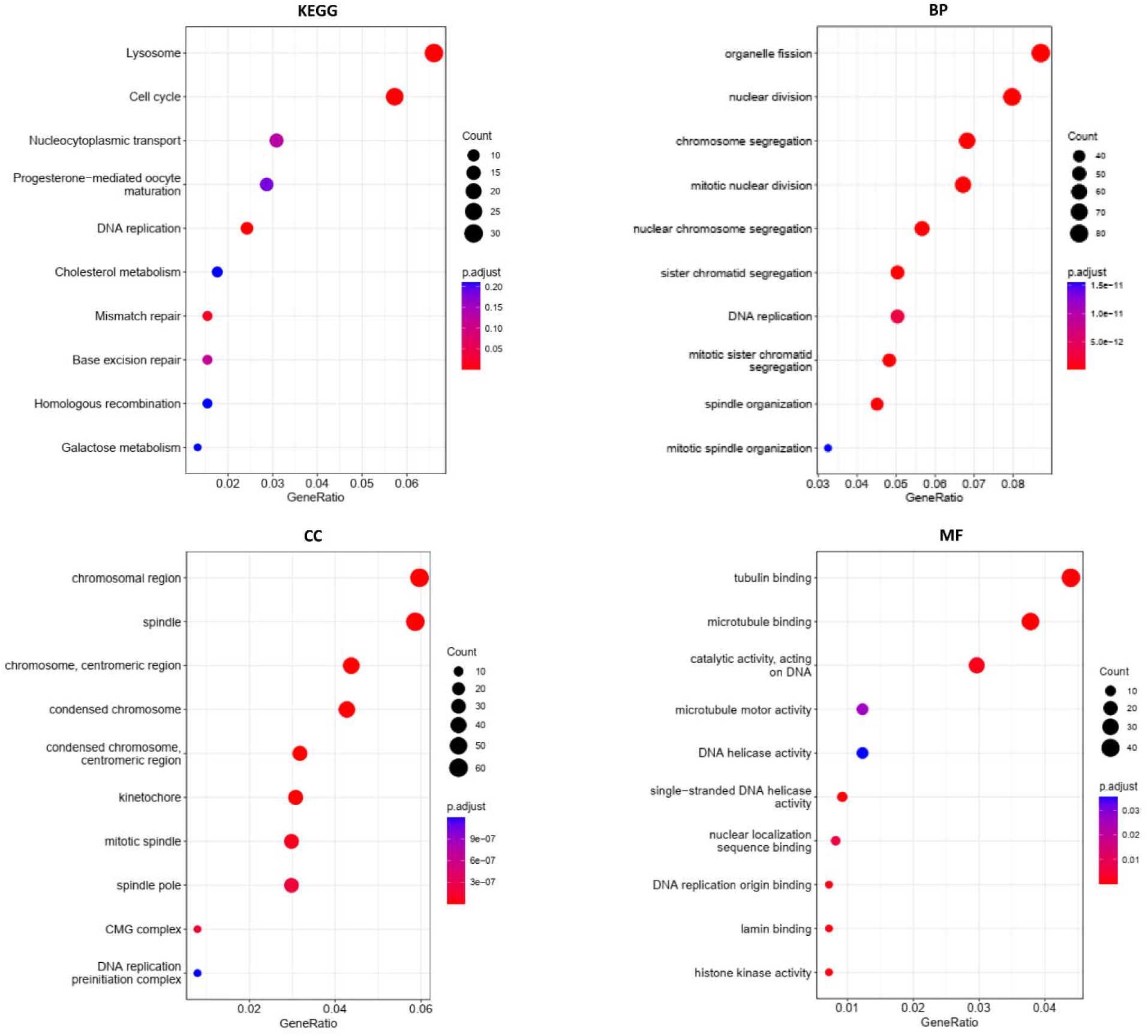
Gene enrichments in MLL-rearranged acute leukemia treated with LAMP-5 shRNAs. KEGG analysis; BP: Biological processes; CC: Cellular components; MF: Molecular functions.

### The protein-protein interaction (PPI) network and Reactome analyses

We constructed the PPI network by using 1040 nodes and 9267 edges with the Cytoscape software (combined score > 0.2). The top ten genes with the highest degree scores were presented in Table 2. By analyzing the PPI network, we further constructed the top two clusters in Figure 3. Finally, we constructed the Reactome map by using the PPI network and DEGs (Figure 4) and we performed the Reactome analysis to further identify the top ten Reactome biological processes, including “Cell Cycle, Mitotic”, “Cell Cycle”, “Cell Cycle Checkpoints”, “DNA strand elongation”, “Mitotic Prometaphase”, “Resolution of Sister Chromatid Cohesion”, “Amplification of signal from unattached kinetochores via a MAD2 inhibitory signal”, “Amplification of signal from the kinetochores”, “Activation of ATR in response to replication stress”, and “M Phase” (Supplemental Table S1).

**Table 2.**
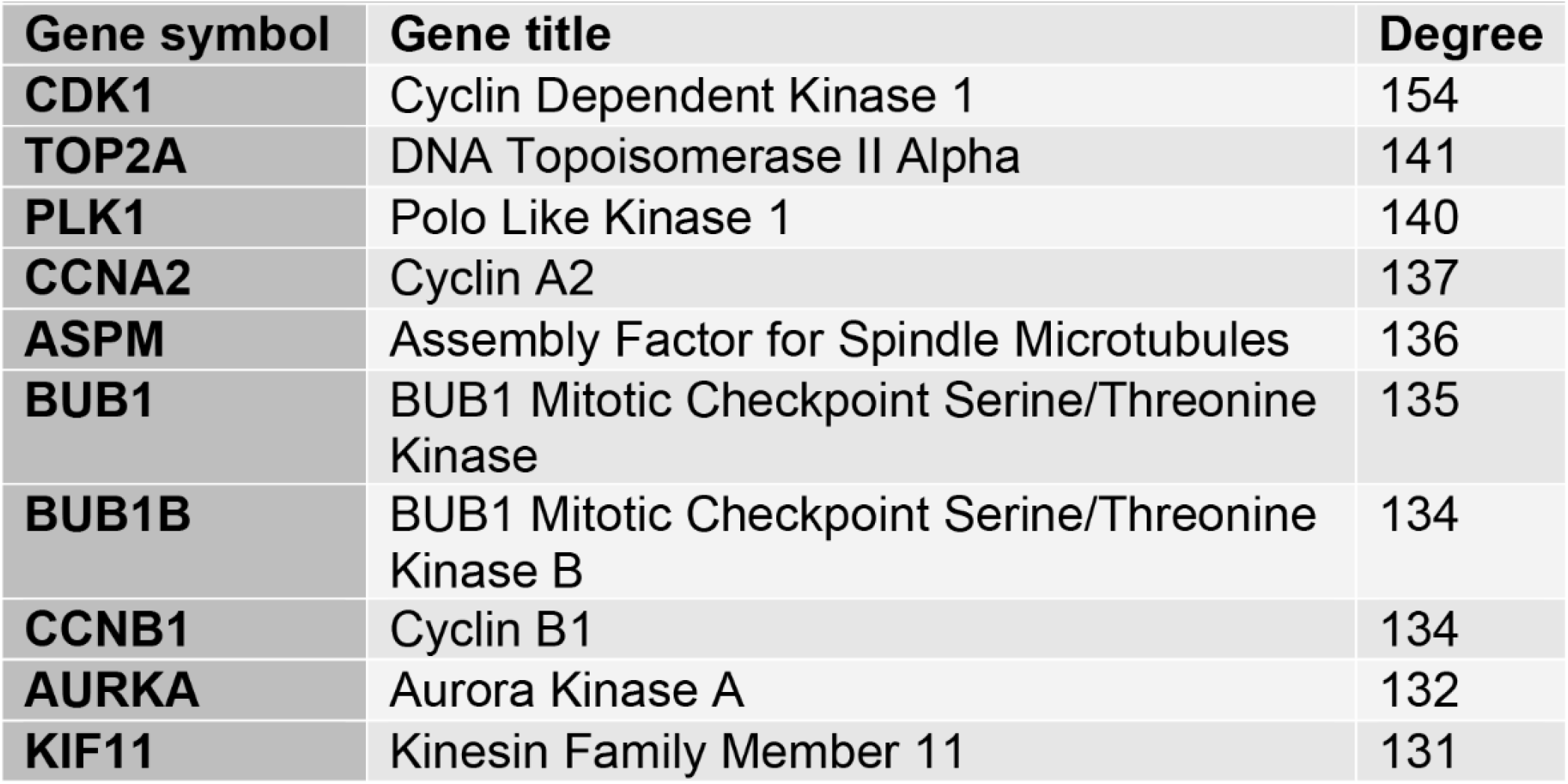
Top ten genes demonstrated by connectivity degree in the PPI network

| **Gene symbol** | **Gene title** | **Degree** |
| --- | --- | --- |
| **CDK1** | Cyclin Dependent Kinase 1 | 154 |
| **TOP2A** | DNA Topoisomerase II Alpha | 141 |
| **PLK1** | Polo Like Kinase 1 | 140 |
| **CCNA2** | Cyclin A2 | 137 |
| **ASPM** | Assembly Factor for Spindle Microtubules | 136 |
| **BUB1** | BUB1 Mitotic Checkpoint Serine/Threonine Kinase | 135 |
| **BUB1B** | BUB1 Mitotic Checkpoint Serine/Threonine Kinase B | 134 |
| **CCNB1** | Cyclin B1 | 134 |
| **AURKA** | Aurora Kinase A | 132 |
| **KIF11** | Kinesin Family Member 11 | 131 |

**Figure 3.**
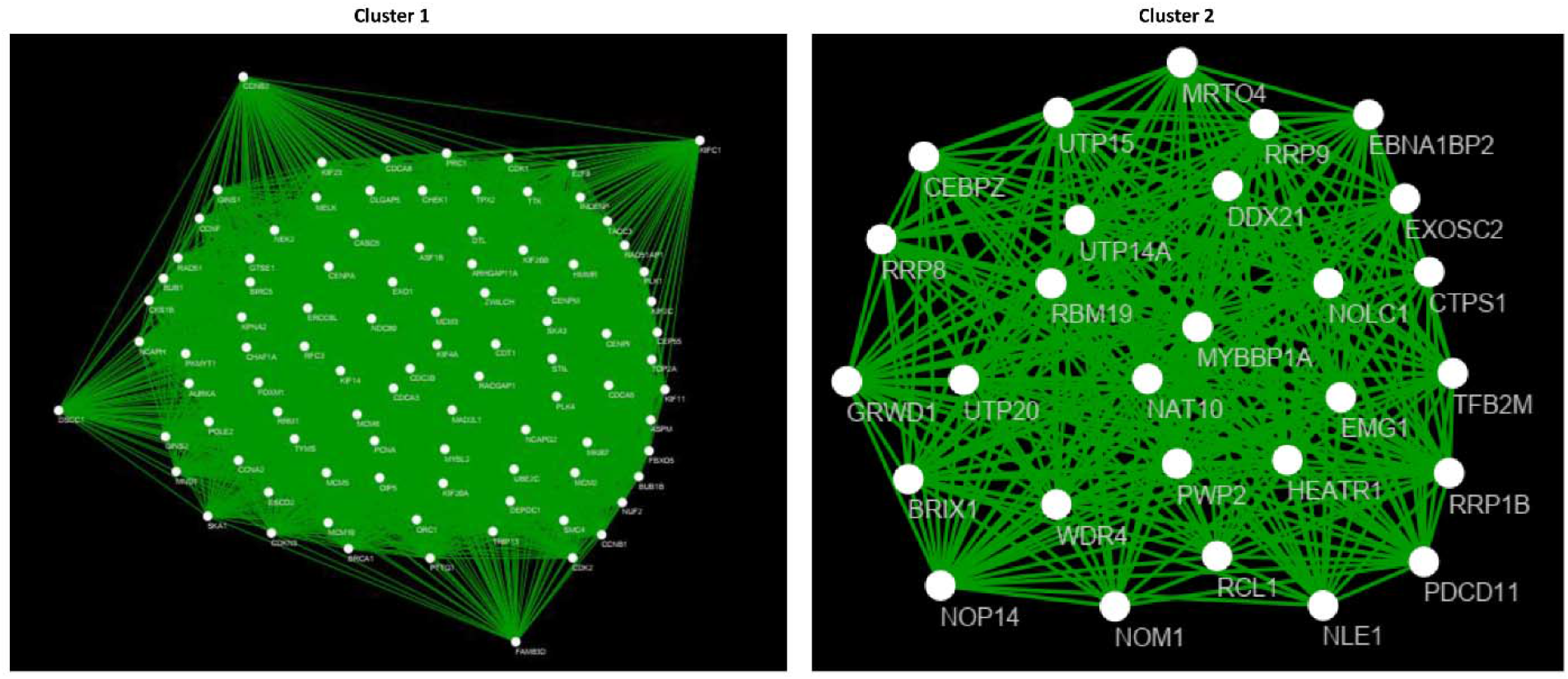
The PPI network analyses of DEGs in MLL-rearranged acute leukemia treated with LAMP-5 shRNAs. The cluster 1 and cluster 2 were constructed by MCODE.

**Figure 4.**
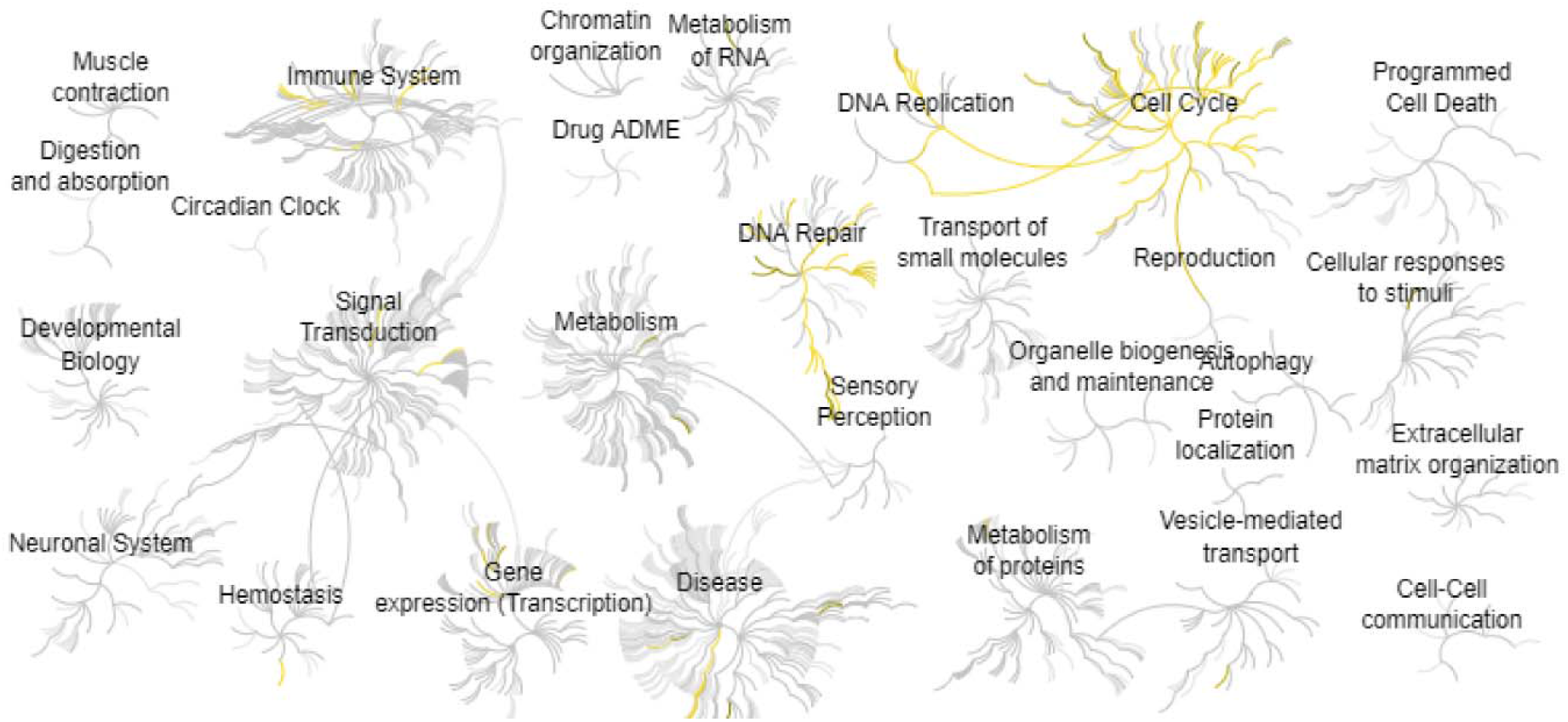
Reactome map representation of the significant biological processes in MLL-rearranged acute leukemia treated with LAMP-5 shRNAs.

## Discussion

The MLL is related to chromosomal translocations in a subtype of acute leukemia, which includes acute lymphoblastic leukemia and myeloid leukemia^11^. Interestingly, LAMP5 is a direct target of the oncogenic MLL-fusion protein, which is a potential target for the treatment of MLL-rearranged acute leukemia^12^. In this study, we obtained the key molecules and signaling pathways by bioinformatic method, which will help us understand the mechanisms of LAMP5-regulated leukemia.

Our study identified the most important signaling pathways “Lysosome” and “Cell Cycle” that strongly correlated with the LAMP5 mediated MLL-rearranged acute leukemia. Similarly, Sreoshee Rafiq et al reported that lysosomes play an important role in degrading cellular macromolecules and recently showed the ability to regulate the tumor microenvironment in leukemia^13^. Interestingly, Karolin Eifler et al also reported that SUMOylation can control the cell cycle in the progression of cancers^14^.

Our study also found the critical molecules by creating the PPI network based on the LAMP5-regulated leukemia data (Table 2). Similarly, Z Huang et al found that CDK1 increases the interaction of SOX2 to further enhance the stemness of lung cancer cells^15^. G protein-coupled receptors are essential for cell functions and are involved in several diseases^16-34^. P M Cobelens et al found that hydrogen peroxide damages GRK2 (GPCR-related protein) via calpain and cdk1-mediated signaling pathway^35^. Fan Zhang et al found that the inhibition of TOP2A by miRNA-599 can repress the malignancy of bladder cancer cells^36^. The study by Lilia Gheghiani et al showed that PLK1 leads to chromosomal instability to promote the progression of cancer^37^. The study by Yichao Wang showed that CCNA2 acts as a potential marker for breast cancer^38^. Qian Yang et al found that ASPM is a key biomarker that is closely related to the cell proliferation of colorectal cancer cells^39^. The study by Li-Jing Zhu et al showed that BUB1 can increase the proliferation of cancer cells by regulating the phosphorylation of SMAD2^40^. The study by Juan-Hua Tian et al showed that BUB1B is critical for cancer cell proliferation and is strongly correlated with poorer outcomes for patients with cancer. Moreover, BUB1B can control MELK which is important for cancer progression^41^. Hui Zhang et al found the CCNB can regulate the cell cycle and aging by mediating the p53 singling pathway in cancers^42^. Ruijuan Du et al reported that AURKA is critical for cancer treatment and found that AURKA can control the functions of AURKA substrates to regulate tumor suppressors or oncogenes^43^. Circadian genes are essential molecules in the cells, which can regulate the gene expression and protein modification to further regulate the cell functions in several diseases such as cancer^44-55^. Cristina Cadenas found that the loss of circadian clock genes can enhance tumor progression by regulating cancer-related genes such as AURKA^56^. Izabela Neska-Długosz et al found the KIF11 gene expression is changed in cancer tissues, which is related to the outcome of patients with colorectal cancers^57^.

In conclusion, this study identified the critical molecules in the MLL-rearranged acute leukemia with the knockdown of LAMP5. The major signaling pathways are “Lysosome” and “Cell Cycle” signaling pathways. Therefore, our study may provide insights into the treatment of leukemia by mediating LAMP5.

## Author Contributions

Zhiwen Qian, Wei Zhang: Methodology, Conceptualization, Writing-Reviewing and Editing.

